# Intrinsic equivalence between developmental prosopagnosia questionnaires

**DOI:** 10.1101/267351

**Authors:** Daisuke Matsuyoshi, Katsumi Watanabe

## Abstract

The 20-item prosopagnosia index (PI20) is a self-report measure of face recognition ability, which is aimed to assess the risk for developmental prosopagnosia (DP), developed by Shah, Gaule, Sowden, Bird, and Cook (2015). Although they validated PI20 in several ways and it may serve as a quick and cost-effective measure for estimating DP risk (Livingston & Shah, in press; Shah et al., 2015), they did not formally evaluate its validity against a pre-existing alternative questionnaire (Kennerknecht et al., 2008) even though they criticized the weak relationship of the pre-existing questionnaire to actual behavioral face recognition performance. Thus, we administered the questionnaires to a large population (N = 855) and found a very strong correlation (r = 0.82 [95% confidence interval: 0.80, 0.84]), a principal component that accounted for more than 90% of the variance, and comparable reliability between the questionnaires. These results show unidimensionality and equivalence between the two questionnaires, or at least, a very strong common latent factor underlying them. The PI20 may not be greater than the pre-existing questionnaire; the two questionnaires measured essentially the same trait. The intrinsic equivalence between the questionnaires necessitates a revision of the view that the PI20 overcomes the weakness of the pre-existing questionnaire. Because both questionnaires contained unreliable items, we suggest, instead of using either questionnaire alone, that selection of a set of items with high reliability may offer a more robust approach to capture face recognition ability.

## 1. Introduction

Prosopagnosia is a neurodevelopmental condition that has deficits in recognizing human faces. Individuals with acquired prosopagnosia have lost their ability in recognizing faces after brain damage, whereas individuals with developmental prosopagnosia (DP) suffer from face recognition at any time during their development, in the absence of any brain injury. DP is characterized by selective deficits in recognizing faces, despite normal intelligence and intact visual object recognition ability (Brad Duchaine & Nakayama, 2005), and is estimated to affect 2% of the population (Kennerknecht, Grueter, et al., 2006; Kennerknecht, Plumpe, Edwards, & Raman, 2006).

It is evident that neurological injury is the primary cause of the inability to recognize faces in the case of acquired prosopagnosia, but what lies behind the DP is still unclear, because distinct anatomical anomalies are not apparent. Despite recent efforts to find neural signatures for DP (Garrido et al., 2009; Rosenthal et al., 2017; Rossion et al., 2003; Song, Zhu, Li, Wang, & Liu, 2015), quantitative, biological measure for estimating DP risk is not established. This absence of confirmed biological markers creates challenges in diagnosing DP by behavioral criteria alone (Barton & Corrow, 2016). It therefore forces us to rest on subjective report for being bad at faces when diagnosing (or more precisely, specifying or classifying) DP. In fact, almost all DP studies have recruited suspected DPs solely based on self-report complaining of face recognition difficulties or semi-structured interviews at best. Although some post-hoc behavioral validation may or may not be employed, there still is no established way to determine whether the suspected DPs represent simply the low-end, quantitatively extreme expression of face recognition ability or qualitatively distinct pathology (see Barton & Corrow, 2016).

Common behavioral testing for DP includes Cambridge Face Memory Test (CFMT) (Brad Duchaine & Nakayama, 2006), Cambridge Face Perception Test (CFPT) (Bradley Duchaine, Germine, & Nakayama, 2007), and famous face tests (Brad Duchaine & Nakayama, 2005; McNeil & Warrington, 1993). In addition to these behavioral tests, questionnaire-based tests are developed to serve as a quick and cost-effective measure for estimating DP risk. Kennerknecht, Ho, and Wong (2008) developed a 15-item DP questionnaire in Hong Kong population (hereafter, HK questionnaire) and estimated the prevalence of DP to be about 2%, which is in the same range as in Caucasians. However, the HK questionnaire has been criticized because it contains items irrelevant to face recognition and it has a weak relationship to actual behavioral performance (Palermo et al., 2017; Shah, Gaule, Sowden, Bird, & Cook, 2015). Recently, Shah et al. (2015) developed a new DP questionnaire, the 20-item prosopagnosia index (PI20), which was aimed to overcome the weakness of the pre-existing questionnaire, and showed that the PI20 correlated with behavioral face recognition performance. In addition, Shah and colleagues claimed that “adults have good insight into their face recognition difficulties” in general populations (Livingston & Shah, in press). However, the PI20 was not formally evaluated against the pre-existing Kennerknecht’s questionnaire and thus the relationship between the questionnaires remains unclear. Given that items in the two questionnaires are so similar simply asking so how good (or bad) people are at recognizing individual faces, it is likely that the two questionnaire measures essentially the same traits. We examined this issue by administering the two questionnaires to a large population and performing a set of analyses including correlation analysis, hierarchical clustering, and a brute-force calculation/comparison of reliability coefficients. If the PI20 is a greater self-report instrument in identifying DP than the pre-existing questionnaire, the PI20 is expected to have a moderate relationship with the HK questionnaire such that items of the two questionnaires form different clusters and/or have higher scale reliability than the HK questionnaire.

## 2. Materials and Methods

### 2.1 Participants

Eight hundred and fifty-five young Japanese adults (427 female, 428 male; mean age: 20.9 ± 2.2 [±1 SD] years; range: 18–36 years) participated in the study. All had normal or corrected-to-normal vision and none reported a history of neurological or developmental disorders. The study procedure was conducted in accordance with the Declaration of Helsinki and approved by the Committee of Ethics, Waseda University, Japan (#2015-033). All participants provided written informed consent prior to participation.

### 2.2 Procedure

We asked participants to complete the questionnaires using an 8-inch touchscreen tablet PC. They were required to indicate the extent to which 36 items (15 from the pre-existing Hong Kong (HK) prosopagnosia questionnaire (Kennerknecht et al., 2008), and 20 from PI20 (Shah et al., 2015), and an additional item pertaining self-confidence in face recognition ability: “I am confident that I can recognize faces well compared to others”) described their face recognition experiences. Responses were provided using a five-point Likert scale ranging from 1 (strongly disagree) to 5 (strongly agree). The participants were instructed to complete the questionnaires at their own pace.

### 2.3 Data analysis

Because the HK prosopagnosia questionnaire developed by Kennerknecht et al. (2008) contains four dummy questions (HK#10, #11, #12, and #13) that are irrelevant with respect to face recognition, we excluded these items and calculated the total scores ranging 11 to 55, using the remaining 11 items (hereafter, ‘HK11’) [score range: 11-55]. PI20 scores were calculated using all 20 items and ranged from 20 to 100. As females have exhibited superior performance in behavioral face recognition studies [3], we examined sex differences between the questionnaire scores. In addition, we used polychoric correlation coefficients to infer latent Pearson correlations between individual items from the ordinal data. The polychoric correlation matrix was estimated using two-step approximation (Olsson, 1979).

Cronbach’s α and Revelle & Zinbarg’s omega total coefficients were calculated to assess the scale reliability of both HK11 and PI20. Omega total coefficients were estimated using a maximum likelihood procedure (Revelle & Zinbarg, 2009). CIs for the coefficients were estimated using a bootstrap procedure (10,000 replications) with a bias-corrected and accelerated approach (DiCiccio & Efron, 1996; Kelley & Pornprasertmanit, 2016). As it was possible that higher reliability coefficients merely reflected the higher number of items in the PI20, relative to that in the HK11 (Cortina, 1993), we performed a brute-force calculation of reliability coefficients for all 167,960 (_20_C_11_) possible combinations of PI20 items, 11 items at a time (i.e., subsets of the PI20 generated by choosing 11 of the 20 items), which allowed us to compare reliability coefficients between the questionnaires with a virtual match of the numbers of items.

## 3. Results

### 3.1 Total scores and score distribution

Table 1 shows descriptive statistics for the total HK11 and PI20 scores. Independent two sample t tests showed no significant differences between in HK11 (*t_853_* = 0.0511, *p* = 0.9592, Cohen’s *d* = 0.0035 [95% confidence interval (CI): −0.1306, 0.1376] or PI20 (*t_810_* = 0.9578, *p* = 0.3384, Cohen’s *d* = 0.0655 [95% CI: −0.0686, 0.1996]) scores between males and females. In addition, Bayesian analysis using a JZS prior (*r* scaling = 1) (Rouder, Speckman, Sun, Morey, & Iverson, 2009) showed strong evidence for the null hypothesis (i.e., no sex difference) for both HK11 (Bayes factor *BF_10_* = 0.0544) and PI20 (*BF_10_* = 0.0856) scores. In addition, two-sample Kolmogorov-Smirnov tests showed no significant sex differences between the distributions (Figure 1) of HK11 (*p* = 0.9982) and PI20 (*p* = 0.7554) scores. These results indicate that females and males showed almost identical mean HK11 and PI20 scores and score distributions, suggesting that sex was not a significant factor.

**Table 1.** Descriptive statistics of total scores for the questionnaires. A higher score indicates lower self-reported face recognition skills. Note that females and males showed similar total scores in terms of not only summary statistics, but also distribution, as shown in Figure 1.

| Questionnaire | Sex | Mean | SD | Min | Median | Max |
| --- | --- | --- | --- | --- | --- | --- |
| HK11 | Female | 24.05 | 6.72 | 11 | 23 | 48 |
|  | Male | 24.03 | 6.72 | 11 | 23 | 49 |
|  | Total | 24.04 | 6.71 | 11 | 23 | 49 |
| PI20 | Female | 48.33 | 13.00 | 26 | 46 | 89 |
|  | Male | 47.49 | 12.70 | 24 | 46 | 85 |
|  | Total | 47.91 | 12.85 | 24 | 46 | 89 |

**Figure 1.**
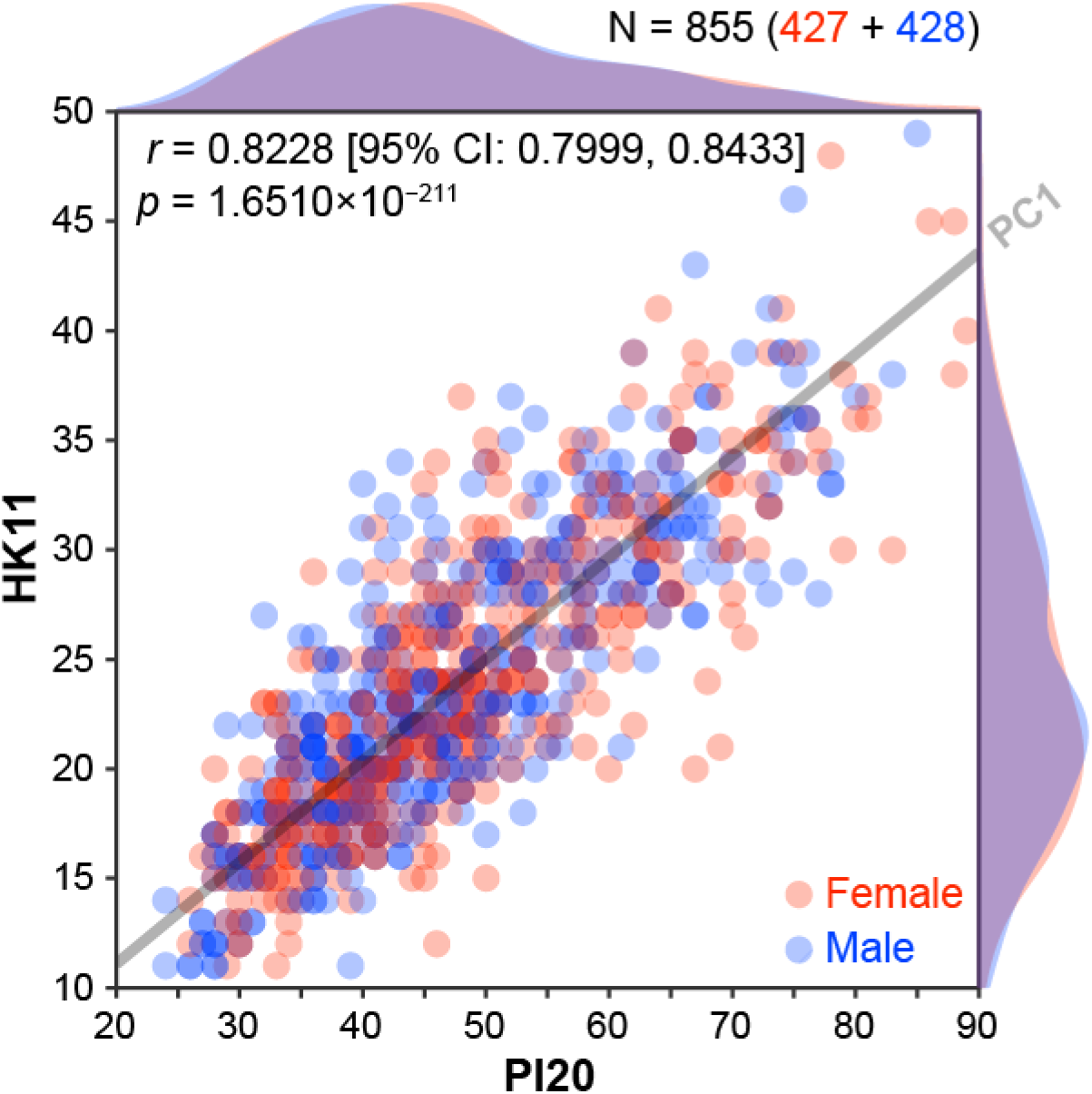
Correlation between total scores for the two prosopagnosia questionnaires. Scatter plot with color-coded transparent density curves of total scores for the 20-item prosopagnosia index (x-axis) and 11 items from Hong Kong prosopagnosia questionnaire (y-axis). Each circle indicates individuals’ data, and color represents sex (red = female scores; blue = male scores). The gray transparent line represents a linear orthogonal regression line (first principal component, PC1 axis), which accounts for more than 90% of the total variance in scores in PCA with a singular value decomposition.

### 3.2 Correlations between total scores

The results showed a very strong significant correlation between the total scores for the two questionnaires (Figure 1, *r* = 0.8228 [95% CI: 0.7999, 0.8433], *p* = 1.6510 × 10^−211^), suggesting a significant overlap of prosopagnosia traits assessed via each measure. It should be noted that Fisher’s z test showed no significant sex difference in the correlation between total scores (difference: *r_diff_* = 0.0065 [95% CI: −0.0371, 0.0502], *z* = 0.2917, *p* = 0.7705; females: *r* = 0.8200 [95% CI: 0.7863, 0.8489], *p* = 4.7087 × 10^−105^; males: *r* = 0.8265 [95% CI: 0.7939, 0.8543], *p* = 2.3806 × 10^−108^). Principal component analysis (PCA) with singular value decomposition of the correlation matrix between total scores showed that the first principal component (PC1) accounted for 91.1% (using standardized scores) and 94.2% (using raw scores) of the total variance in scores.

### 3.3 Correlations between individual item scores

The correlation matrix (Figure 2) generally showed correlations between individual items *across* the two scales; however, some items were not correlated with other items to the extent that they would reduce the reliability or internal consistency of a single measure pertaining to a single construct (i.e., DP risk). In fact, hierarchical clustering using the unweighted pair group method with arithmetic mean showed that eight out of 36 items were distant from a cluster to which most items belonged (shaded areas in Figure 2, dendrogram). These eight items consisted of (Table 2): the four items already known to be irrelevant with respect to DP (HK#10, HK#11, HK#12, and HK#13), and two items from the HK prosopagnosia questionnaire (HK#2 and HK#7), and two items from the PI20 (PI#3 and PI#13). Previous studies reported that five of the eight item-score differences were marginal (score difference [DP − control]: <1) between individuals with suspected DP and typically developed control individuals (0.45 for HK#10, −0.39 for HK#11, 0.11 for HK#12, −0.45 for HK#13, and 0.62 for PI#3) (Kennerknecht et al., 2008; Shah et al., 2015). However, it should be noted that the score difference exceeded 1 for the remaining three items (1.46 for HK#2, 1.12 for HK#7, and 1.16 for PI#13), suggesting that these three items could actually measure prosopagnosia traits that differ from those measured via the other 28 items.

**Table 2.** Test items shown with the mean scores. Shaded items are distant from a cluster to which the bulk of the items belonged (see Figure 2). Items marked with asterisks (*) are reverse scored.

| Test item |  | Mean score<br>(1 SD) |  |
| --- | --- | --- | --- |
|  |  | Females | Males |
| HK#1 * | I can easily follow actors in a movie | 2.94<br>(1.21) | 2.83<br>(1.14) |
| HK#2 | Some of my family have problems in recognizing faces | 1.49<br>(0.96) | 1.48<br>(0.89) |
| HK#3 | People often tell me I do not recognize them | 2.48<br>(1.19) | 2.49<br>(1.21) |
| HK#4 * | I can decide immediately if a face is familiar | 2.48<br>(1.08) | 2.41<br>(1.08) |
| HK#5 | It takes me a long time to recognize people | 2.83<br>(1.14) | 2.74<br>(1.14) |
| HK#6 * | I always recognize family members | 1.15<br>(0.53) | 1.15<br>(0.58) |
| HK#7 * | I can easily form a mental picture of a red rose | 1.65<br>(0.86) | 1.95<br>(1.06) |
| HK#8 * | I can easily form pictures of close friends in my mind | 1.56<br>(0.87) | 1.60<br>(0.91) |
| HK#9 * | I recognize famous people immediately | 2.59<br>(1.19) | 2.59<br>(1.15) |
| HK#10 * | I can decide immediately whether a face is male or female | 1.93<br>(0.92) | 1.87<br>(0.78) |
| HK#11 | I get lost in new places | 3.77<br>(1.24) | 3.07<br>(1.28) |
| HK#12 * | I can see if a face is attractive | 2.51<br>(1.08) | 2.42<br>(1.07) |
| HK#13 | I have problems reading emotions in a face | 2.42<br>(1.02) | 2.53<br>(1.07) |
| HK#14 | I avoid meetings as I might overlook familiar people | 1.79<br>(0.94) | 1.86<br>(0.99) |
| HK#15 | I do not recognize people the day after a brief meeting | 3.09<br>(1.32) | 2.93<br>(1.32) |
| CONF * | I am confident that I can recognize faces well compared to others | 3.33<br>(1.16) | 3.16<br>(1.22) |
| PI#1 | My face recognition ability is worse than most people | 2.82<br>(1.09) | 2.68<br>(1.10) |
| PI#2 | I have always had a bad memory for faces | 2.11<br>(1.08) | 2.06<br>(1.02) |
| PI#3 | I find it notably easier to recognize people who have distinctive facial features | 4.23<br>(0.78) | 4.29<br>(0.78) |
| PI#4 | I often mistake people I have met before for strangers | 2.49<br>(1.17) | 2.51<br>(1.22) |
| PI#5 | When I was at school I struggled to recognize my classmates | 2.05<br>(1.11) | 1.92<br>(1.02) |
| PI#6 | When people change their hairstyle, or wear hats, I have problems recognizing them | 2.69<br>(1.16) | 2.71<br>(1.16) |
| PI#7 | I sometimes have to warn new people I meet that I am 'bad with faces' | 1.76<br>(1.13) | 1.72<br>(1.09) |
| PI#8 * | I find it easy to picture individual faces in my mind | 2.45<br>(1.12) | 2.31<br>(1.01) |
| PI#9 * | I am better than most people at putting a 'name to a face' | 3.47<br>(1.15) | 3.25<br>(1.19) |
| PI#10 | Without hearing people's voices, I struggle to recognize them | 2.06<br>(1.01) | 2.15<br>(1.02) |
| PI#11 | Anxiety about face recognition has led me to avoid certain social or professional situations | 1.86<br>(1.08) | 1.86<br>(1.04) |
| PI#12 | I have to try harder than other people to memorize faces | 2.33<br>(1.18) | 2.34<br>(1.19) |
| PI#13 * | I am very confident in my ability to recognize myself in photographs | 1.84<br>(0.95) | 1.97<br>(1.00) |
| PI#14 | I sometimes find movies hard to follow because of difficulties recognizing characters | 2.23<br>(1.19) | 2.08<br>(1.11) |
| PI#15 | My friends and family think I have bad face recognition or bad face memory | 1.79<br>(1.04) | 1.76<br>(0.98) |
| PI#16 | I feel like I frequently offend people by not recognizing who they are | 1.75<br>(0.97) | 1.71<br>(0.96) |
| PI#17 * | It is easy for me to recognize individuals in situations that require people to wear similar clothes (e.g. suits, uniforms and swimwear) | 2.67<br>(1.13) | 2.53<br>(1.14) |
| PI#18 | At family gatherings, I sometimes confuse individual family members | 1.66<br>(0.97) | 1.63<br>(0.94) |
| PI#19 * | I find it easy to recognize celebrities in 'before-they-were-famous' photos, even if they have changed considerably | 3.44<br>(1.07) | 3.46<br>(1.06) |
| PI#20 | It is hard to recognize familiar people when I meet them out of context (e.g. meeting a work colleague unexpectedly while shopping) | 2.63<br>(1.23) | 2.53<br>(1.23) |

**Figure 2.**
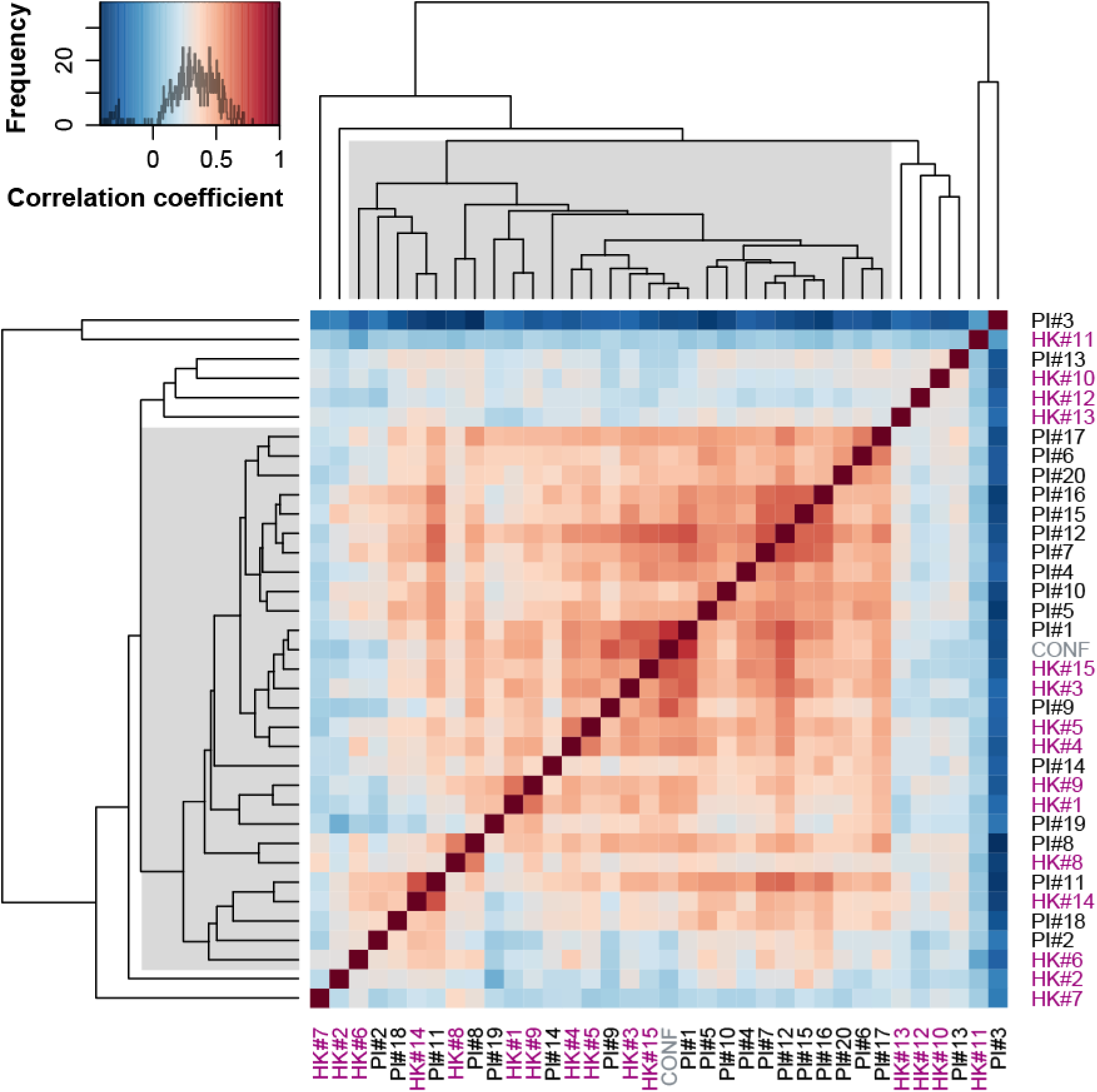
Polychoric correlation matrix and hierarchical clustering (dendrogram) for individual item scores. Polychoric correlation coefficients are color-coded using the color key shown at the top left histogram. The dendrogram was obtained by a hierarchical clustering based on Pearson correlation distances using the unweighted pair group method with arithmetic mean. CONF, the question pertaining self-confidence in face recognition ability.

### 3.4 Scale reliability

We found that the reliability coefficients for the PI20 were higher relative to those for the HK11 (HK11: *α* = 0.8449 [95% CI: 0.8273, 0.8633], *ω_t_* = 0.8767 [95% CI: 0.8571, 0.8880]; PI20: *α* = 0.9174 [95% CI: 0.9102, 0.9249], *ω_t_* = 0.9368 [95% CI: 0.9300, 0.9424]). Follow-up Feldt paired tests (Feldt, 1980) confirmed significant differences in reliability coefficient between the HK11 and PI20 (difference in *α*: *t_853_* = 16.4437, *p* = 5.5132 × 10^−53^; difference in *ω_t_*: *t_853_* = 17.4868, *p* = 9.4696 × 10^−59^).

The brute-force calculation of reliability coefficients showed that the coefficients for the eleven-item PI20 subsets were almost comparable (within 1 SD) to those for the HK11 (*α*: mean = 0.8530 ± 0.0392 [±1 SD], median = 0.8474, range = 0.7495–0.9324; *ω_t_*: mean = 0.8914 ± 0.0225 [±1 SD], median = 0.8933, range = 0.8122–0.9438). This indicated that the HK11 and PI20 demonstrated almost equivalent reliability at the individual-item level.

## 4. Discussion

Our results showed that the two representative ‘DP questionnaires’ are closely related to each other such that they are almost indistinguishable in terms of correlation analyses, PCA, hierarchical clustering, and item reliability. It is worth noting that a recent meta-analysis showed that test-retest reliabilities for instantaneously administered tests are about *r* = 0.8 (Calamia, Markon, & Tranel, 2013), which is comparable to our findings (*r* = 0.82). This may indicate that the correlation coefficient between the two questionnaires is sufficiently high to consider that the two questionnaires might measure essentially the same trait to the extent of reliability that solid neuropsychological tests can achieve. PCA also confirmed that the primary component accounted for more than 90% of the total variance.

Shah et al. (2015) claimed that “the PI20 can play a valuable role in identifying DP” and it will lead to “reliable classification [of DP]”; however, the huge correlation and comparable reliability between the two questionnaires indicate that the PI20 is not at all greater than the pre-existing questionnaire, even though they criticized it because of its “weak relationship” to actual behavioral face recognition performance. Although further research is needed to evaluate/compare the PI20 and HK11 with behavioral face recognition performance, it may be hard to expect that, given the huge correlation, it is only the PI20 that has a strong relationship to face recognition performance whereas the HK11 has a weak relationship to it. In fact, recent studies suggest that people have poor insight into their face recognition abilities (Bowles et al., 2009; Palermo et al., 2017).

In addition, we found strong correlations between individual items across the two questionnaires. Instead of forming different clusters of their own, the bulk of the items across two questionnaires formed a single cluster, which is distant from the remaining eight items (Table 2). These eight items consisted of four items related to face *processing*, but not pertaining to their own face *recognition* abilities (the presence of face recognition difficulties in their family [HK#2]; ability to judge facial gender [HK#10], facial attractiveness [HK#12], and facial emotion [HK#13]), two items related to face recognition but not correlated with most other items (ability to remember individuals with distinctive facial features [PI#3]; confidence in finding themselves in photographs [PI#13]), and two items not related to face recognition (navigation deficits [HK#11]; ability to mentally image an object [HK#7]). Although we did not administer other questionnaires, the little or no correlation between face-recognition-relevant and irrelevant items supports the view that the high inter-individual correlations may be specific to self-reported face recognition ability. In terms of scale reliability, we suggest researchers and clinicians to discard these eight items when assessing an individual’s insight into their face recognition ability.

In conclusion, our results indicated that the two representative DP questionnaires (Kennerknecht et al., 2008; Shah et al., 2015) essentially measured the same face recognition traits. In addition, the huge correlation and robust principal component demonstrated a common latent factor between the two measures. This putative unidimensionality could be intrinsic to insight into one’s face recognition ability, rather than behavioral face recognition performance per se (Palermo et al., 2017). Although the PI20 may serve as a measure for estimating DP risk and face recognition ability in general populations (Livingston & Shah, in press), our findings suggest that, contrary to the Shah et al.’s claims, its reliability and validity may be almost equivalent to that of the pre-existing questionnaire (Kennerknecht et al., 2008) (precisely, the reduced subset, HK11) at the individual-item level. Given the current state of DP, where neither objective diagnostic criteria nor biological markers have been established (Barton & Corrow, 2016; Susilo & Duchaine, 2013), it might be a good idea to focus on creating a reliable face recognition questionnaire (rather than a ‘DP questionnaire’) that can predict behavioral face recognition performance by discarding unreliable, face-recognition-irrelevant items (Table 2, but see Palermo et al., 2017; Zell & Krizan, 2014). Alternatively, more exploratory research not only using HK11 and PI20 together or a combination thereof, but also a range of other face processing measures could aid the extraction of latent prosopagnosia traits/dimensions and the development of a valid DP taxonomy.

## Acknowledgement

The datasets and code are available at https://github.com/dicemt/matsuyoshi2018dp (doi: 10.5281/zenodo.1189186).

## Authors’ contributions

DM and KW designed the study and wrote the manuscript. DM collected and analyzed the data.

